# Behavioral and transcriptomic analyses of mecp2 function in zebrafish

**DOI:** 10.1101/2023.09.13.557635

**Authors:** Nicholas J. Santistevan, Colby T. Ford, Cole S. Gilsdorf, Yevgenya Grinblat

**Affiliations:** University of Wisconsin – Madison, Departments of Integrative Biology and Neuroscience, Madison, WI, USA; University of Wisconsin – Madison, Genetics Ph.D. Training Program, Madison, WI, USA; University of North Carolina at Charlotte, School of Data Science, Charlotte, NC, USA; University of North Carolina at Charlotte, Department of Bioinformatics and Genomics, Charlotte, NC, USA; Tuple LLC, Charlotte, NC, USA

**Author notes:** Correspondence (NJS); (YG).

**Keywords:** MeCP2, zebrafish, transcriptomic analysis, Rett Syndrome (RTT), behavioral analysis

## Abstract

Rett Syndrome (RTT), a human neurodevelopmental disorder characterized by severe cognitive and motor impairments, is caused by dysfunction of the conserved transcriptional regulator Methyl-CpG-binding protein 2 (MECP2). Genetic analyses in mouse Mecp2 mutants, which exhibit key features of human RTT, have been essential for deciphering the mechanisms of MeCP2 function; nonetheless, our understanding of these complex mechanisms is incomplete. Zebrafish mecp2 mutants exhibit mild behavioral deficits but have not been analyzed in depth. Here we combine transcriptomic and behavioral assays to assess baseline and stimulus-evoked motor responses and sensory filtering in zebrafish mecp2 mutants from 5-7 days post-fertilization (dpf). We show that zebrafish mecp2 function is dispensable for gross movement, acoustic startle response, and sensory filtering (habituation and sensorimotor gating), and reveal a previously unknown role for mecp2 in behavioral responses to visual stimuli. RNA-seq analysis identified a large gene set that requires mecp2 function for correct transcription at 4 dpf, and pathway analysis revealed several pathways that require MeCP2 function in both zebrafish and mammals. These findings show that MeCP2’s function as a transcriptional regulator is conserved across vertebrates and supports using zebrafish to complement mouse modeling in elucidating these conserved mechanisms.

## Introduction

*Methyl-CpG-binding protein 2* (*MECP2*) is an X-linked, transcriptional regulatory gene associated with a number of disorders marked by cognitive impairment, including autism, schizophrenia, and attention-deficit hyperactivity disorder (1–3). Mutations in *MECP2* are causally linked to Rett Syndrome (RTT), a neurodevelopmental disorder that affects 1∼ in 10,000-20,000 girls and is characterized by cognitive defects, impaired motor coordination, loss of language skills and hand use, and seizures (4–6). Patients with RTT exhibit normal postnatal development for the first 6-18 months of life, followed by developmental stagnation and a rapid regression of motor and cognitive function (5). RTT patients present with reductions in brain volume, neuronal soma size and dendritic arborization, and immature synaptic connections (7).

Animal models of MeCP2 dysfunction have been critical to understanding the role of MeCP2 in neurodevelopment, cognitive and motor function, and transcriptional regulation. MeCP2 is conserved across vertebrates, with the transcriptional regulatory (TRD) and methyl-binding domains (MBD) sharing strong amino acid identity between humans, mice, and zebrafish (8). In mice, brain-specific deletions of *Mecp2* cause neuronal aberrations similar to those observed in RTT patients, including impaired synaptic physiology, impaired dendritic morphogenesis (9, 10), and motor and cognitive deficits typical of RTT (11–17). Similarly, *mecp2^Q63X^* homozygous zebrafish, which harbor a predicted loss-of-function mutation, exhibit decreased thigmotactic responses, decreased activity, and mild motor dysfunction in response to mechanical stimulation (18).

Zebrafish is a powerful model system for exploring how MeCP2 functions in the brain to control distinct aspects of behavior. *mecp2* is expressed broadly in the zebrafish embryo, including the brain, as early as 24-48 hours post-fertilization (hpf) (8, 19). Its expression at this early time point is notable as the circuits that mediate several stereotyped behaviors develop between 24 and 96 hours and begin to function immediately thereafter (20, 21). Here we used a two-pronged approach to ask how *mecp2* functions in zebrafish: behavioral assessment and transcriptomic (RNA-seq) analysis. Using a panel of assays to quantify the initiation, execution, and modulation of several baseline and stimulus-evoked behaviors, we show largely normal baseline behaviors and normal measures of sensory filtering (acoustic habituation and prepulse inhibition (PPI)), in 5 days post-fertilization (dpf) *mecp2* mutant zebrafish. Notably, *mecp2* mutant zebrafish showed an aberrant response to visual stimuli that has not been reported in mouse mutants.

In many tissues and molecular pathways, MeCP2 functions as a methyl-binding transcription factor that acts as both a transcriptional activator and a repressor (22, 23). Our RNA-Seq analysis of 4 dpf *mecp2* mutants identified a large set of genes that require *mecp2* function for correct transcription, and pathway analysis showed enrichment of several pathways previously associated with Rett syndrome, indicative of a deep conservation of MeCP2 functions in vertebrates. Collectively, these results suggest that zebrafish is a useful, yet under-utilized model system that could provide valuable insights into the function of MeCP2, contributing to our understanding of neurodevelopmental disorders associated with MeCP2 dysfunction.

## Results

### mecp2 mutant zebrafish show normal spontaneous locomotion

Patients with RTT and mouse models of MeCP2 dysfunction exhibit impaired mobility. To evaluate motor function in *mecp2^Q63X^* homozygous mutant zebrafish, which harbor a predicted loss-of-function mutation (18), we assessed 5-7 dpf larval zebrafish for baseline activity levels, kinematics, and their responses to various stimulus paradigms. We first quantified baseline activity in *mecp2* mutant larvae by measuring gross movement over time and the frequency of spontaneous initiation of a turn or swim maneuver. To do this, we recorded 6 dpf *mecp2* mutant larvae and wild-type siblings for 3 minutes and measured several aspects of their swimming behaviors, namely, distance moved, initiations of turns, and initiations of swimming bouts. Over the 3-minute recording period, *mecp2* mutant and wild-type siblings showed similar gross movement with no difference in the distance swam (Fig. 1A). Next, we divided the 3-minute recording window into 30 second intervals and assessed larvae for turn initiations and swimming bouts during each interval. Across the recording window, *mecp2* mutant and wild-type siblings exhibited similar probability of initiating turns (Fig. 1B; 6 recording intervals for both genotypes) and swimming bouts (Fig. 1C; 6 recording intervals for both genotypes). Together, these results indicate that 6 dpf *mecp2* mutants show normal baseline activity and normal locomotion. *mecp2* mutants show normal baseline activity and normal locomotion.

**Fig. 1.**
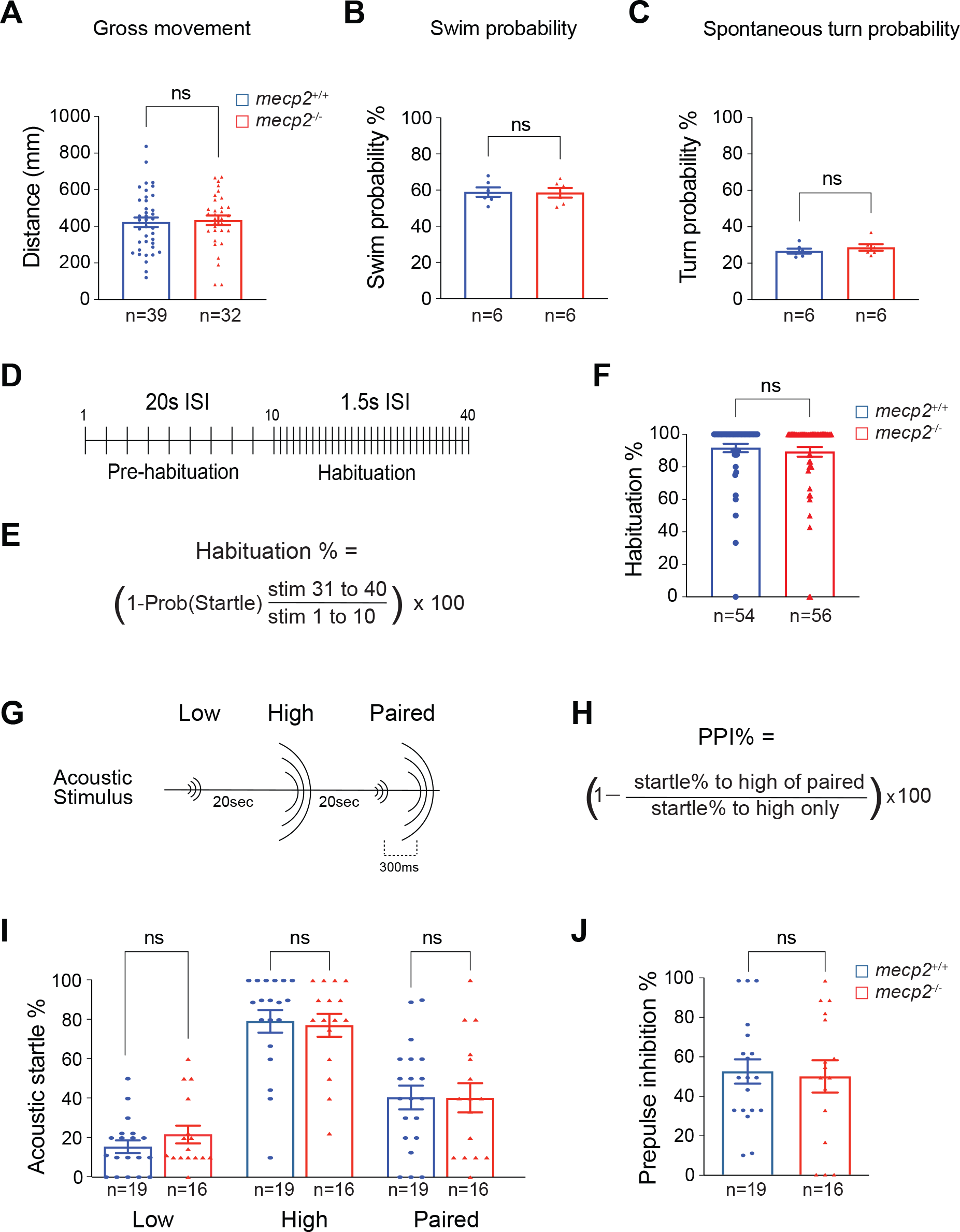
*mecp2* mutants exhibit normal baseline activity and measures of sensory filtering. (A-C) Baseline motor activity levels between *mecp2* mutant larvae and wild-type siblings including overall gross movement (A), probability of initiating swimming bouts (B), and probability of initiating spontaneous turns (C). (D, E) Schematic representation of the habituation assay (D) and formula used to calculate habituation (E). Stimuli 1-10 were delivered at non-habituating 20s intervals and stimuli 11-40 were delivered at habituating 1.5s intervals. (F) Mean habituation of wild-type (shown in blue) and *mecp2* mutant (shown in red) larvae. Schematic representation of the prepulse inhibition (PPI) assay (G) and formula used to calculate PPI% (H). (I) Startle responsiveness of *mecp2* mutants and wild-type siblings to low, high, and paired acoustic stimuli. (J) Mean PPI% of *mecp2* mutants and wild-type siblings. Number of larvae shown at the bottom of the graphs. *p<0.05, **p<0.01,Mann-Whitney test versus wild-type. Mann-Whitney. Error bars denote SEM.

### mecp2 mutants show normal sensory filtering of the acoustic startle response

Since loss of Mecp2 function in mice results in impaired acoustic startle response (ASR) (11, 24), we asked if loss of mecp2 function impaired acoustic startle behavior in larval zebrafish. By 5 dpf, zebrafish have established the neural circuits sufficient to control ASRs with kinematic parameters similar to those observed in adult zebrafish (25). The ASR is driven by the Mauthner hindbrain reticulospinal interneuron, the command neuron of the ASR (26). Each Mauthner neuron receives direct acoustic input from sensory hair cells via the VIIIth nerve (27). The Mauthner sends descending commissural axons down the spinal cord to stimulate contralaterally positioned motor neurons, which induces a contralateral trunk muscle contraction and results in the zebrafish turning away from the acoustic stimulus (26, 28–30). To examine the initiation, performance, and modulation of ASR behavior in *mecp2* larvae, we used a semi-automated behavioral platform that delivers a series of acoustic stimuli at defined intensities and intervals, records ASRs with a high-speed camera, and tracks individual larvae’s movement to quantify aspects of ASR behavior.

The circuit that drives the ASR in zebrafish exhibits experience-dependent modulation of the ASR, namely, habituation learning and sensorimotor gating (20). Habituation has been termed the simplest form of non-associative learning that is exhibited by all animals (31) and is observed as a progressive decline in responsiveness to repeated, insignificant stimuli. To explore the requirement for *mecp2* function in ASR modulation, we assessed habituation by administering 10 high-intensity acoustic stimuli at non-habituating intervals of 20 seconds, followed by 30 high-intensity acoustic stimuli at habituating intervals of 1.5s (Fig. 1D). Habituation was calculated by taking the ratio of responses to the habituating and non-habituating stimuli (Fig. 1E) (29). Analysis of habituation between *mecp2* mutant larvae and wild-type siblings revealed that *mecp2* mutant zebrafish habituate normally (Fig. 1F).

Next, we measured prepulse inhibition (PPI) of the ASR (20), a widely used measure of sensorimotor gating in animal models and humans (32), thought to specifically reflect the function of inhibitory microcircuits (33). PPI of the ASR measures the ability of a weak acoustic stimulus to attenuate an animal’s responsiveness to a subsequently presented, strong acoustic stimulus (34). Larvae were exposed to a series of acoustic stimuli, including low intensity stimuli that rarely evoke an ASR, high intensity stimuli that typically cause an ASR, and paired stimuli to measure PPI, in which a low intensity stimulus was delivered 300 milliseconds preceding a high intensity stimulus (Fig. 1G). PPI percentage was calculated by taking the ratio of responses to the high of the paired stimuli over the responses to the high stimulus alone (Fig. 1H) (20). At each intensity level, acoustic startle was the same in *mecp2* mutants and wild-type siblings (Fig. 1I) indicative of normal PPI in *mecp2* mutants (Figure 1J).

### mecp2 mutant zebrafish exhibit aberrant visually-guided behaviors

The visual system is emerging as a reliable biomarker of MeCP2 disruption with both mouse models and RTT patients displaying lowered visual acuity (35, 36). Conditional Mecp2 deletion in GABAergic parvalbumin-expressing cells has been shown to abolish experience-dependent critical period plasticity of visual circuits and impair visual acuity (37). To ask if mecp2 function is required for visual behaviors in zebrafish, we compared the initiation and execution of stereotyped visually-evoked behaviors in 5-7 dpf *mecp2* mutants. It is important to note these responses are not mediated by the Mauthner neuron and show kinematics that are distinct from the ASRs (25). Following adaptation to a given light intensity, zebrafish larvae perform a turning maneuver when exposed to a sudden increase in light (‘light flash’) or the abrupt absence of light (‘dark flash’) (25, 38, 39). Following exposure to light flashes, 7 dpf *mecp2* mutants showed an increased response frequency compared to wild-type siblings (Fig. 2A). Analysis of the kinematic properties of light flash evoked turns showed that *mecp2* mutants execute turn responses with a shorter mean initiation latency, i.e. faster (Fig. 2B). The mean angle turned while executing the light flash evoked turn was greater in *mecp2* mutants (Fig. 2C). The mean duration of the light flash evoked turn was also greater in *mecp2* mutants (Fig. 2D) but mean swim velocity was indistinguishable from wild-type siblings (Fig. 2E).

**Fig. 2.**
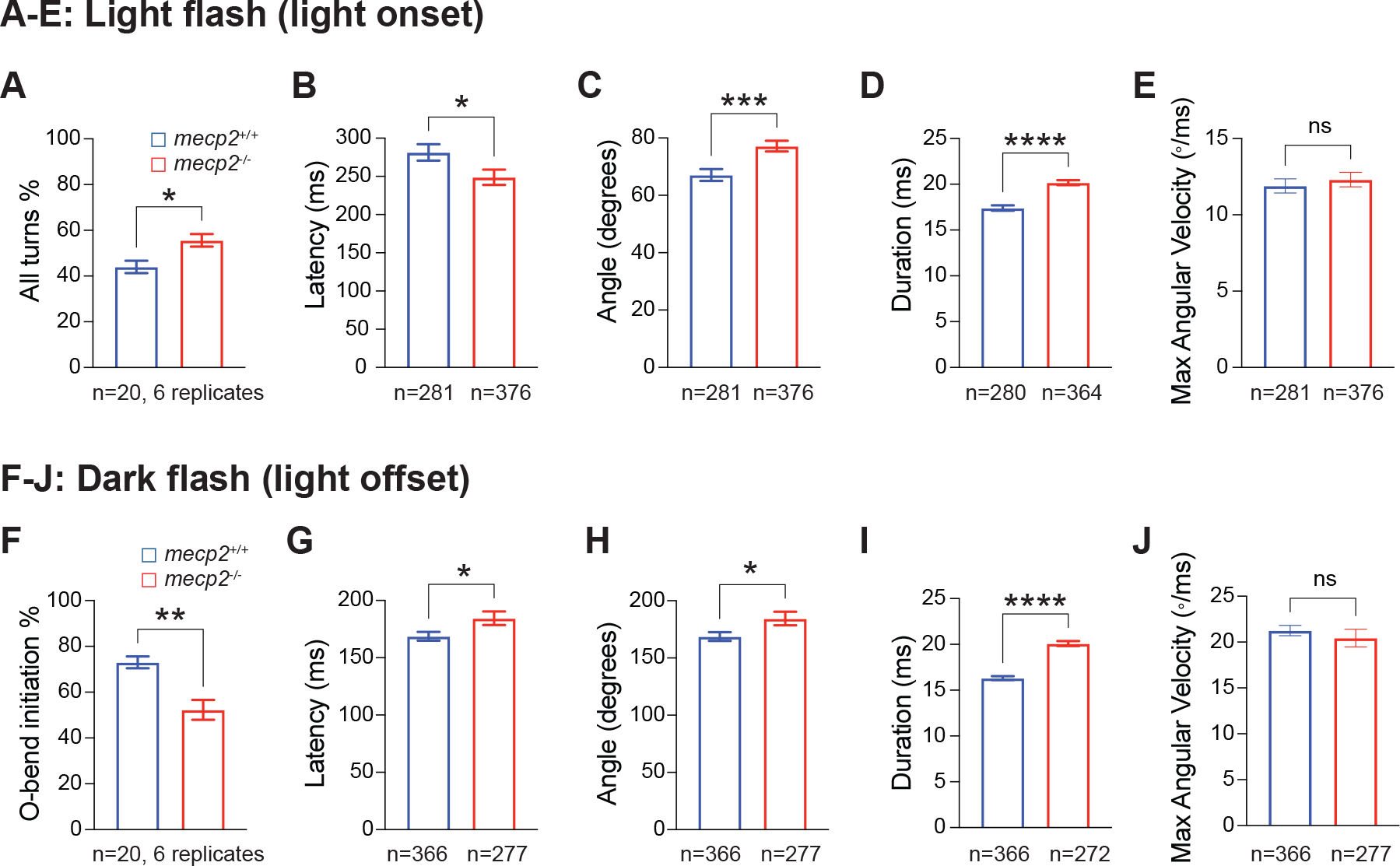
*mecp2* mutants exhibit aberrant kinematic parameters and responses to light offset and light onset. (A) Light flash elicits a low angle turn response. Mean frequency of turns initiated to a series of 10 light flash stimuli. wild-type siblings are shown in blue and *mecp2* mutants are shown in red. *p<0.05, Mann-Whitney test versus wild-type. Number of larvae across 6 replicates shown at the bottom of graph. Error bars denote SEM. (B-E) Kinematic analysis of the light flash induced turning response including turn latency (B), turn angle (C), turn duration (D), and turn angular velocity (E). *p<0.05, ***p<0.001, ****p<0.0001. Mann-Whitney test versus wild-type. Number of responses measured shown at the bottom of graphs. Error bars denote SEM. (F) Dark flash elicits O-bend response. Mean frequency of O-bend responses initiated to a series of dark flash stimuli. wild-type siblings are shown in blue and *mecp2* mutants are shown in red. **p<0.01, Mann-Whitney test versus wild-type. Number of larvae across 6 replicates shown at the bottom of graph. Error bars denote SEM. (G-J) Kinematic analysis of the dark flash induced O-bend response including turn latency (G), turn angle (H), turn duration (I), and turn angular velocity (J). *p<0.05, ****p<0.0001, Mann-Whitney test versus wild-type. Number of responses measured shown at the bottom of graphs. Error bars denote SEM.

In response to an abrupt absence of light, or “dark flash”, larval zebrafish execute an O-bend, characterized by a large angle turn in which they reorient their bodies by 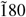 degrees (25, 38). We evaluated the probability of 5 dpf *mecp2* mutants to initiate an O-bend turn maneuver to dark flashes after the larvae had adapted to light and found a marked reduction in O-bend initiations in response to dark flashes (Fig 2F). Kinematic analysis of O-bend initiation showed an increase in mean latency to initiate the O-bend in mutants compared to wild-types (Fig 2G). The mean angle turned while performing the O-bend was also increased in *mecp2* mutants (Fig 2H) and the mean duration of the O-bend turn was markedly longer in *mecp2* mutants compared to wild-types at all dark flash intensities (Fig 2I) while mean swim velocity was indistinguishable between wild-types and mutants (Fig. 2J). Together, these results suggest a role for *mecp2* in mediating the detection or response to the light OFF and ON retinal pathways.

### Transcriptomic analysis reveals conserved RTT-associated pathways in zebrafish

Next, we performed bulk RNA-Seq transcriptomic analysis comparing *mecp2* mutant larvae and wild-type siblings at 4 dpf, the earliest stage when the circuit that drives the acoustic startle response is both fully developed and fully functional. RNA was prepared from pools of 3 individual *mecp2* mutant embryos and pools of 3 age-matched wild-type embryos, with 10 replicates per genotype, reverse-transcribed and sequenced on the Illumina platform. Bioinformatic analysis, carried out as described in Materials and Methods, identified 2659 differentially expressed genes (DEGs) in *mecp2* mutant homozygotes (Benjamini-Hochberg corrected FDR <0.05); transcript levels were significantly up-regulated at 1,394 loci and down-regulated at 1,265 loci (Fig. 3A, Supplemental Table 1, see Supplementary Data in the GitHub repository).

**Fig. 3.**
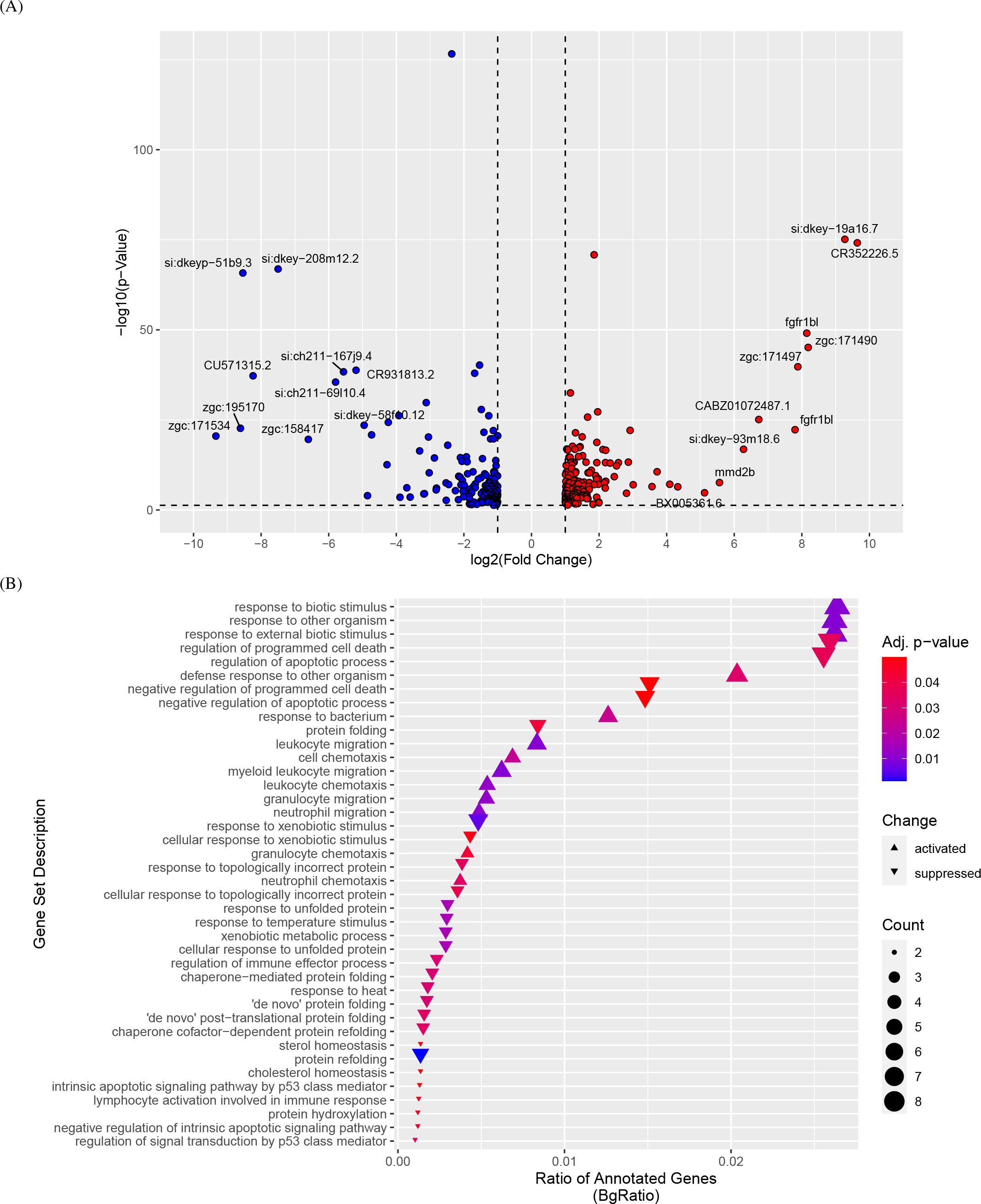
Transcriptome analysis identifies activated and suppressed genes and gene sets in *mecp2* mutants. (A) A volcano plot depicting the fold change and statistical significance of individual genes, with log_2_ (Fold Change) by -log_10_ (p-value) of individual genes shown as points and colorized by their significance and change in regulation (grey is non-significant, red is significant and upregulated, blue is significant and downregulated). The names of top 10 upregulated genes and top 10 downregulated genes by log_2_ (Fold Change) were labeled. (B) The top 40 differentially expressed gene sets in *mecp2* mutants relative to wild-type siblings, identified by GSEA, with 14 activated and 26 suppressed gene sets with adj. *p*-value < 0.05. This plot shows the BgRatio (the ratio of annotated genes) by gene set descriptor, points colored by the adjusted *p*-value, sized by count of genes in each set, and points shaped by the overall change (activated vs. suppressed).

We examined the DEGs for expression of two known MeCP2 target genes, NMDA receptors *bdnf* (40), and *creb1* (41), and found them unchanged in *mecp2* mutants. Expression levels of MeCP2 homologs *mbd1, mbd2, mbd3,* and *mbd4* were likewise either unchanged or decreased in *mecp2* mutants (*mbd1a*, fold decrease= -0.236, p value = 0.007) (Supplemental Table 1, see Supplementary Data in the GitHub repository).

Kyoto Encyclopedia of Genes and Genomes (KEGG) analysis and Gene Set Enrichment Analysis (GSEA) are bioinformatic tools used to analyze gene expression data and identify enriched biological pathways. KEGG is a comprehensive knowledge base that provides information on gene functions and molecular networks by mapping gene expression data onto the KEGG pathways to identify those that are significantly enriched with differentially expressed genes (42, 43). KEGG analysis of the DEG set revealed 7 pathways significantly affected in *mecp2* mutants, including the G2-M DNA damage checkpoint of the cell cycle and cholesterol homeostasis, among others (adjusted *p* value <0.05) (Table 1). Each of the 7 pathways identified in *mecp2* mutant zebrafish have also been reported in studies of Mecp2-deficient mouse models and/or patients with RTT (see Table 1 for citations). GSEA is a computational method that assesses whether a set of genes, defined by a biological pathway or a functional category, is enriched at the top or bottom of a ranked list of genes (44). We performed GSEA on the expression data and identified 14 molecular pathways that were activated and 26 that were suppressed (Fig. 3B) in the *mecp2* larvae (adjusted p value < 0.05). Among these, cholesterol homeostasis, which was suppressed in *mecp2* mutants, was of note as it was also identified in KEGG analysis and it has been demonstrated that cholesterol homeostasis is disrupted in *Mecp2* mutant mice (45) and is altered in cells derived from RTT patients (46).

**Table 1.**
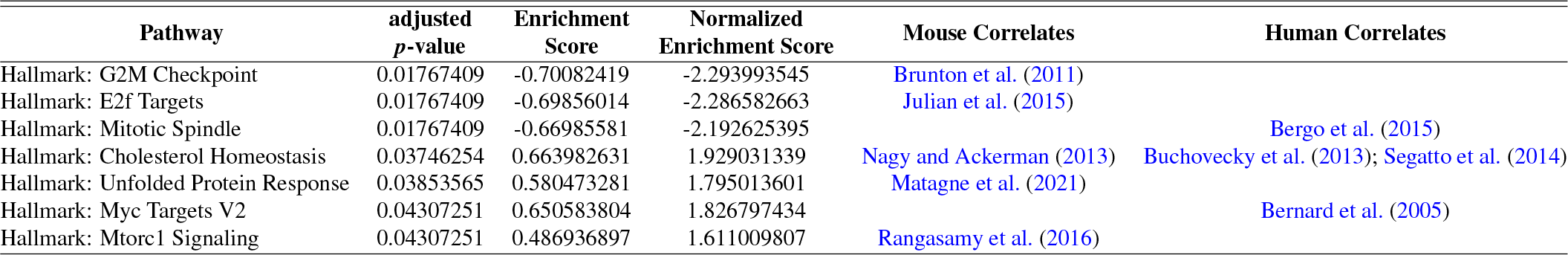
Pathway analysis reveals conserved pathways between zebrafish, MECP2 mouse models, and RTT patients. KEGG analysis revealed several pathways affected by loss of *mecp2* in 4 dpf larval zebrafish. All pathways are conserved between *mecp2* mutant zebrafish, Mecp2-deficient mouse models, and/or patients with RTT. Pathways with adjusted *p* values <0.05 are shown. The full list of leading-edge genes with altered expression for each of these pathways is listed in Supplemental Table 2, see Supplementary Data in the GitHub repository.

| Pathway | adjusted *p*-value | Enrichment Score | Normalized Enrichment Score | Mouse Correlates | Human Correlates |
| --- | --- | --- | --- | --- | --- |
| Hallmark: G2M Checkpoint | 0.01767409 | -0.70082419 | -2.293993545 | Brunton et al. (2011) |  |
| Hallmark: E2f Targets | 0.01767409 | -0.69856014 | -2.286582663 | Julian et al. (2015) |  |
| Hallmark: Mitotic Spindle | 0.01767409 | -0.66985581 | -2.192625395 |  | Bergo et al. (2015) |
| Hallmark: Cholesterol Homeostasis | 0.03746254 | 0.663982631 | 1.929031339 | Nagy and Ackerman (2013) | Buchovecky et al. (2013); Segatto et al. (2014) |
| Hallmark: Unfolded Protein Response | 0.03853565 | 0.580473281 | 1.795013601 | Matagne et al. (2021) |  |
| Hallmark: Myc Targets V2 | 0.04307251 | 0.650583804 | 1.826797434 |  | Bernard et al. (2005) |
| Hallmark: Mtorc1 Signaling | 0.04307251 | 0.486936897 | 1.611009807 | Rangasamy et al. (2016) |  |

Previous work in mice has shown that loss of Mecp2 function yields increased brain cholesterol concentration which has detrimental effects on neuron development and function due to perturbed cholesterol homeostasis (45, 50). The mevalonate pathway is responsible for the production of isoprenoids, which are essential for various cellular functions, including cholesterol synthesis (54), while the steroid biosynthesis pathway is involved in the catabolism and elimination of cholesterol (55). The mevalonate and steroid biosynthesis pathways work together to maintain cholesterol homeostasis in the body. Tracing the mevalonate-cholesterol pathway through defined KEGG pathways (dre:00900, dre:00100, and dre:00140) and overlaying mutant vs. wild-type expression of each gene exemplifies the changes in expression due to *mecp2* mutation (56) (Fig. 4). Specifically, both the mevalonate pathway and steroid biosynthesis pathways are activated in *mecp2* mutant zebrafish, leading to altered cholesterol synthesis.

**Fig. 4.**
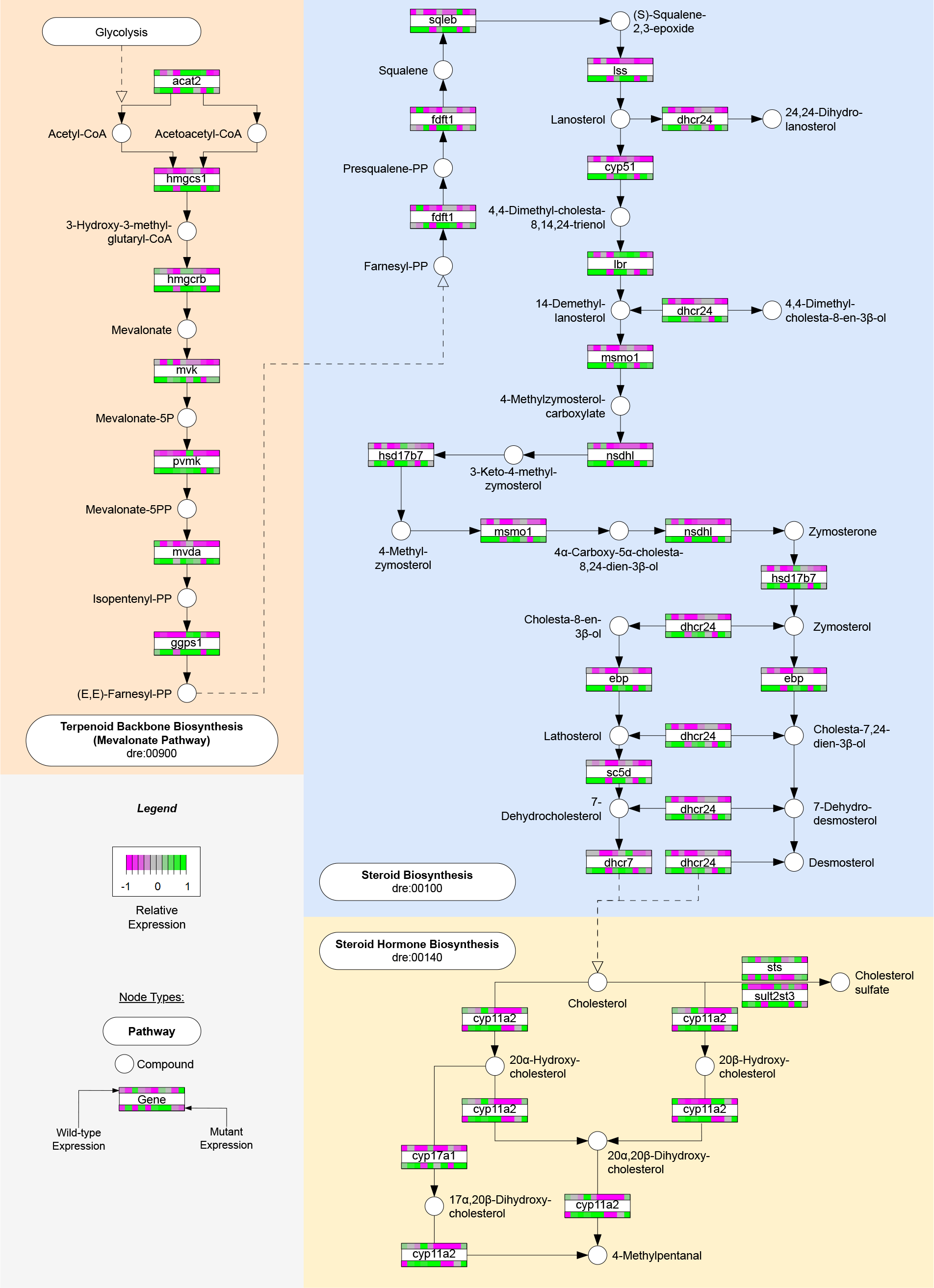
Relative gene expression differences shown between *mecp2* mutants and wild-type siblings throughout the mevalonate-cholesterol pathway. A combined mevalonate-cholesterol pathway, simplified from KEGG pathways dre:00900, dre:00100, and dre:00140 to show only genes relevant to this study and *D. rerio*. Color bars on each gene node represent the relative gene expression of each sample in this study, scaled between [-1,1]. The top color bar of a given gene node represents the 10 wild-type samples and the bottom color bar represents the 10 mutant samples.

## Discussion

Impairments in sensorimotor gating of the ASR have been documented in Mecp2-deficient mouse models and in patients with RTT, and we asked if this aspect of MeCP2 function was conserved in a teleost model, the zebrafish. Using a panel of behavioral assays in *mecp2* mutants aged 5-7 days, we show that mecp2 function is dispensable for ASR, but is required for correct visually-guided startle responses in zebrafish. Genome-wide transcriptional analysis of *mecp2* mutants identified a large set of differentially expressed genes and several molecular pathways that require MeCP2 function in zebrafish and in mammals; these include glycolysis, the G2-M DNA damage checkpoint, and cholesterol homeostasis. This conservation, together with the unique advantages of zebrafish for high-throughput drug screening and long-term behavioral analysis, support the utility of the zebrafish model system for dissecting conserved mechanisms of MeCP2 function in vertebrates.

### Modeling motor deficits associated with Rett Syndrome

Previous research has shown that mice with brain-specific deletions of Mecp2 exhibit impaired synaptic physiology and motor and cognitive deficits typical of RTT (9– 11, 16, 57). These deficits include limited mobility, decreased motor coordination measured using rotarod assays, and decreased cognition measured using cued and conditional tasks, with deficits associated with learning and memory (13–15, 58, 59). A prior study found that *mecp2* mutation in zebrafish results in increased spontaneous contractions at 25 hpf and an increased duration of head tactile-evoked startle responses at 51 hpf (18), which may be indicative of an excitation/inhibition imbalance in *mecp2* mutants. This study also reported that loss-of-function mutation of *mecp2* in zebrafish larvae at 6 dpf leads to reduced thigmotaxis, decreased activity, and swimming bouts with lower velocity (18). Interestingly, morpholino-mediated knockdown of *mecp2* at 28 hpf and 72 hpf results in decreased motor activity, as measured by slower touch responses and abnormal/slower swimming, respectively (19). Our assessment of locomotor activity in 6 dpf *mecp2* mutant zebrafish indicate that *mecp2* function is dispensable for additional measures of baseline motor activity, as evidenced by normal overall gross movement, frequency of spontaneous swimming bouts, and frequency of spontaneous turns. Taken together, these analyses provide a comprehensive characterization of baseline activity in free-swimming 6 dpf *mecp2* mutant larvae.

### Modeling cognitive deficits associated with Rett Syndrome

Impaired sensorimotor gating of the acoustic startle response has been observed in conditional *Mecp2* knockout mice (11) and disorders associated with MECP2 dysfunction (60–63). Unexpectedly, we found no impairments in experience-dependent modulation of the ASR in *mecp2* mutants, as both habituation learning and sensorimotor gating, measured by PPI, were normal. Importantly, there are discrepancies in how sensory filtering deficits manifest in humans with RTT and Mecp2-null mouse models. In RTT patients, sensorimotor gating is often reduced (32, 64), while in *Mecp2* knockout mice (both global and GABA neuron-specific) increased PPI has been observed (11, 65).

The discrepancy in PPI between the zebrafish and murine Mecp2 loss-of-function models may reflect differences in how PPI is measured in these models and/or the presence of additional behavioral deficits in the *Mecp2* knockout mice that could influence their PPI measures. In mice, the ASR is measured as a graded response to stimulus intensity, measured by the downward force applied to a testing platform when startled (66). PPI is evident when the force generated during an ASR to an intense stimulus preceded by a weak stimulus is less than that generated to an intense stimulus alone. In *Mecp2* knockout mice, reduced acoustic sensitivity and motor deficits (Chao et al., 2010; Ure et al., 2016) can generally diminish the ASR to high-intensity stimuli, indirectly leading to an enhanced PPI phenotype. While the zebrafish ASR shares kinematic and pharmacodynamic properties with mammalian ASR (20, 67), the zebrafish ASR is an all-or-nothing behavior, defined by the initiation of a “C-bend”. PPI in zebrafish is indicated by a reduced probability of C-bend initiation to an intense stimulus that is preceded by a weak stimulus, compared to the probability of C-bend initiation to a high stimulus alone (Figs. 3D-3E) (20). Thus, PPI in zebrafish represents suppressed initiation while in mice it reflects a reduction in response intensity. Although the ASR circuitry in zebrafish is highly analogous to the mammalian ASR circuit, albeit simpler, there are likely differences that determine the all-or-nothing versus graded nature of their respective ASR and thus, may explain why PPI is altered in mice and humans, but not zebrafish, following MeCP2 dysfunction. Our study provides insights into the behavioral effects of *mecp2* mutation in zebrafish, highlighting potential species-specific differences in the function of MeCP2 on sensory filtering.

### Modeling visually-guided behaviors

The visual processing and visual memory deficits observed in individuals with RTT (68), combined with the reduced visual acuity in both mouse models and RTT patients (35, 36), highlight the importance of assessing and addressing visual functioning in intervention strategies for RTT. Visual acuity measures are gaining recognition as reliable biomarkers of MeCP2 dysfunction (35, 36) and our study provides the first evidence that mecp2 plays a similar role in the zebrafish model. *mecp2* mutant zebrafish exhibit an enhanced light flash (light onset) response with distinct alterations in kinematic parameters and attenuated responses to dark flash (light offset) stimuli, accompanied by impaired kinematic parameters. These data show that mecp2 is important for the visual system that responds to changes in illumination, but do not distinguish between a requirement for light perception (visual acuity) and motor response circuitry. It is important to note these responses are not mediated by the Mauthner neuron and show kinematics that are distinct from the ASR (20).

Retinal photoreceptors are a likely site for mecp2 function in the visual system as these cells are able to distinguish between light offset and light onset stimuli. Their responses to these stimuli are driven by parallel circuits within the retina that are dependent on the modulation of glutamate release in cone-to-bipolar cell synapses (69). Increased synaptic vesicle release by cones activates OFF bipolar cells in response to light offset stimuli, while reduced synaptic vesicle release by cones activates ON bipolar cells in response to light onset stimuli (38). Taken together, these data suggest that analysis of the structure and function of the synapses between cones and discrete ON/OFF bipolar cell types will further elucidate the role of mecp2 in the visual circuit.

Prior research has indicated that light offset responses rely on insulin-like growth factor (IGF) signaling, which depends on proper cleavage of IGF binding proteins (IGFBPs) by proteases, enabling IGF to bind its receptor (Miller et al., 2018). Our transcriptomic analysis of *mecp2* mutant zebrafish revealed elevated expression of several IGFBP genes, suggesting reduced IGF availability and signaling (Supplemental Table 1, see Supplementary Data in the GitHub repository). Notably, IGF-1 has emerged as a promising therapeutic target for RTT patients, as exogenous IGF-1 supplementation has shown to ameliorate many RTT-like symptoms in *Mecp2* mutant mice (10, 70, 71). Taken together, these results highlight conserved functions of MeCP2 in the visual response pathway and indicate that MeCP2 dysfunction results in a disruption in detection and/or interpretation of changes in illumination, which may be due to dysregulation of photoreceptor synaptic development.

### Molecular pathways regulated by mecp2

A previous study used transcriptomic analysis of zebrafish *mecp2* mutants to demonstrate enrichment of processes relevant to RTT phenotypes; however, this analysis was performed on 6-hpf early gastrula embryos (72) and well before the critical stages of neural circuit formation (24-96 hpf) in the zebrafish embryo (20, 21). We reasoned that RNAseq at 96 hpf (4 dpf) would capture *mecp2* targets that were most relevant to the circuits tested for function in our behavioral assays. Our analysis uncovered 2659 differentially expressed genes and identified multiple molecular pathways with distinct patterns of activation and suppression, including those involved in glycolysis, G2-M DNA damage checkpoint, and cholesterol homeostasis. Among these pathways, cholesterol homeostasis was of particular note since cholesterol metabolism alterations have been documented in RTT patients (46) and even minor disruptions of cholesterol metabolism in mice profoundly impact neuronal development and lead to neurological abnormalities, including impaired limb clasping and tremors (45, 50). Importantly, dysregulation of cholesterol homeostasis was also evident in the brain of *Mecp2*-deficient mice, and the administration of cholesterol-lowering statin drugs improved motor phenotypes and overall health and extended lifespan in *Mecp2* mutant mice (50). Collectively, these data support the exciting possibility that MeCP2 plays a conserved role in regulating cholesterol metabolism, offering a promising avenue for potential therapeutics targeting this pathway in addressing the impact of MeCP2 dysfunction on neurological function and development.

The unexpectedly mild phenotypes exhibited by *mecp2* mutants may be due to genetic compensation of MeCP2 homologs through transcriptional adaptation, a compensatory mechanism that increases transcription of genes that share sequence similarity with a mutated gene (73, 74). While our transcriptome analysis of *mecp2* mutant zebrafish did not produce evidence of transcriptional upregulation of *mecp2* homologs *mbd1, mbd2, mbd3,* and *mbd4* at 4 dpf, it is possible that genetic compensation ameliorates the consequences of mecp2 dysfunction at other stages of development. Symptoms of RTT typically emerge in patients between 6-18 months of age, following a period of normal development marked by achievement of developmental milestones such as crawling or walking (5, 16). The decline in patients is progressive, with motor and cognitive symptoms manifesting in stages over time. Similarly, *Mecp2*-deficient mice exhibit normal development for the first 4 weeks, followed by a progressive deterioration of motor function (Vashi & Justice, 2019), resulting in impaired locomotion and premature death (61). By 7 weeks of age, mutant mice display enhanced PPI that persists into adulthood (57). Despite their behavioral and transcriptomic deficits, homozygous *mecp2* mutant zebrafish appear to develop normally to adulthood, exhibiting no overt morphological defects and maintaining the ability to reproduce (18). In this study, we focused on 4-7 dpf larval zebrafish as these timepoints are considered to be post-developmental for the acoustic startle circuit. It is possible that in zebrafish, as in mammals, the relevant circuits in *mecp2* mutant zebrafish undergo normal development before the onset of motor defects that are mild enough to allow survival in controlled laboratory environments. Behavioral analysis of juvenile and adult zebrafish *mecp2* mutants will test this hypothesis in future studies.

### Zebrafish as a tool for unraveling mechanisms of MeCP2 function

Our transcriptomic data, together with previous proteomic analysis of larval and adult *mecp2* functions (75), demonstrate remarkable conservation of molecular pathways that depend on mecp2 function in zebrafish, mouse and RTT patients. Zebrafish offer a unique advantage for drug treatment studies, as compounds can be readily added to the media, making it a powerful tool for exploring candidate therapeutics. Recent work has demonstrated that regulators of MeCP2 stability can be targeted by drug treatment (76) and these compounds hold significant promise for exploration in the zebrafish model system. Furthermore, adult viability of *mecp2* mutant zebrafish allows behavioral analysis throughout their lifespan, enabling monitoring of progressive declines in motor responses. Collectively, these data provide compelling justification, and serve as a necessary foundation for future in-depth functional analyses of mecp2 in the zebrafish model. Ease of high-throughput behavioral and pharmacological analyses in the zebrafish model would complement and inform mouse studies, aiding in unraveling the complexities of MeCP2 function and designing therapeutic interventions for RTT and related neurodevelopmental disorders.

## Methods

### Generation and Maintenance of Zebrafish

Zebrafish (*Danio rerio*) were maintained according to established methods (77). The *mecp2^Q63X^* line was obtained from Dr. Cecilia Moens via the Zebrafish International Resource Center (ZIRC). All experimental protocols using zebrafish were approved by the University of Wisconsin Animal Care and Use Committee and carried out in accordance with the institutional animal care protocols. Embryos were raised in E3 media at 28°C on a 14 hr/10 hr light/dark cycle through 7 dpf as previously described (78, 79). Embryos were generated from heterozygous crosses of adults carrying the *mecp2^Q63X^* allele.

### Genotyping

Genotyping of the *mecp2^Q63X^* allele is based on the dCAPs assay (80) using the following primers: *mecp2^Q63X^* forward:

5’- ACAGCTGACTATAATACAGACCTGTCAAA -3’

and *mecp2^Q63X^* reverse:

5’- TGGGTTCAGACTTCCCTGCCTCTGCCTGCA -3’.

To genotype the *mecp2^Q63X^* allele, a mismatch SNP (marked in red) has been introduced into the reverse primer. During PCR, this mismatch creates a PstI (New England Biolabs, #R3140) restriction enzyme site in the amplified product derived from the wild-type DNA template. The PstI site is not present in the PCR product containing the *mecp2^Q63X^* mutation.

### Behavioral Analyses

On the day of behavioral testing, the larvae were held in 60mm-wide Petri dishes with 20 larvae in 10mL E3, kept on a white light box for at least 1 hour, and then transferred to a 4x4 (habituation), 6x6 grid (PPI) or 60mm dishes (gross movement, swim probability, turn probability, light flash, and dark flash). Acoustic startle behavior was elicited using an automated behavioral platform in which the intensity and timing of acoustic stimuli could be controlled (20, 39). Acoustic startle responses were elicited with a minishaker (Brüel & Kjær, Model 4810). The acoustic stimuli were of 3 ms duration, with 1000 Hz waveforms. Light offset-induced (dark flash) O-bends and light onset-induced (light flash) turns were elicited and analyzed as previously described (20, 39). Behavioral responses were captured at 1000 frames per second with a MotionPro Y4 video camera (Integrated Design Tools) with a 50 mm macro lens (Sigma Corporation of America) at 512 x 512 pixel resolution.

FLOTE was used to analyze startle responses in an experimenter-independent, automated manner (20). FLOTE tracked the position of individual larvae frame by frame and characterizes locomotor maneuvers (e.g. C-bend, O-bend, routine turn, swim, etc.) according to predefined kinematic parameters that distinguish these maneuvers. To evaluate habituation and measure startle responsiveness and kinematic parameters, 10 high-intensity stimuli were presented at non-habituating interstimulus intervals (ISIs) of 20 seconds followed by 30 high-intensity stimuli presented at 1.5 second ISIs. The degree to which larvae habituate was calculated as previously defined (29, 39). To evaluate PPI, low and high stimuli were presented at a 300 ms ISI and the degree to which larvae demonstrate startle prepulse inhibition was calculated as previously defined (20). For habituation and PPI behavior, we report data representing the Mauthner neuron-dependent, short-latency C-bend startle (SLC) response. For kinematic analysis of ASRs, we report both SLCs and LLCs and their kinematic parameters. For light onset (light flash) responses, we report initiation of all turns and their kinematic parameters. For light offset (dark flash) responses, we report O-bends and their kinematic parameters.

### RNA-Seq Transcriptome Analysis

Embryos derived from incrosses of *mecp2^Q63X^* homozygous adults or wild-type adults were pooled in groups of three and processed for RNA extraction at 4 dpf as previously described (81). cDNA libraries were prepared using the TruSeq stranded mRNA library preparation protocol with poly-A selection and sequenced on the Illumina HiSeq. 2500. 10 biological replicates of each genotype, confirmed from raw sequencing data alignments using the Integrative Genomics Viewer, were analyzed.

Bioinformatic analysis of transcriptomic data adhere to recommended ENCODE guidelines and best practices for RNA-Seq (82). Alignment of adapter-trimmed (83) (Skewer v0.1.123) 2x150 (paired-end; PE) bp strand-specific Illumina reads to the *Danio rerio* GRCz11 genome (assembly accession GCF_000002035.6) was achieved with the Spliced Transcripts Alignment to a Reference (STAR v2.5.3a) software (84), a splice-junction aware aligner, using annotation modifications (v4.3.2) made by (85). Expression estimation was performed with RSEM v1.3.0 (RNASeq by Expectation Maximization) (86). To test for differential gene expression among individual group contrasts, expected read counts obtained from RSEM were used as input into edgeR (v3.16.5) (87). Inter-sample normalization was achieved among each pair of samples with trimmed mean of M-values (TMM) (88). Statistical significance of the negative-binomial regression test was adjusted with a Benjamini-Hochberg FDR correction at the 5% level (89). Prior to statistical analysis with edgeR, independent filtering was performed, requiring a threshold of at least 1 transcript count per million in at least 2 samples, ignoring any prior group assignment. The validity of the Benjamini-Hochberg FDR multiple testing procedure was evaluated by inspection of the uncorrected p-value distribution.

### Expression Visualization and Gene Set Enrichment

A volcano plot was generated to depict the fold change and statistical significance of individual genes, with log_2_(Fold Change) by -log_10_(p-value) of individual genes shown as points and colorized by their significance and change in regulation (grey is non-significant, red is significant and upregulated, blue is significant and downregulated). The names of top 10 upregulated genes and top 10 downregulated genes by log_2_(Fold Change) were labeled (Fig. 3A).

Gene set enrichment analysis (GSEA) was performed using the clusterProfiler v4.6.2 R (90, 91) package. Gene sets were defined using the “drerio_gene_ensembl” dataset from Ensembl.org in the biomaRt v2.54.1 R package (92, 93). A scatterplot of significant gene sets was generated to show the change in expression of the top 40 sets, based on adj. p-value. This plot shows the BgRatio (the ratio of annotated genes) by gene set descriptor, points colored by the adjusted *p*-value, sized by count of genes in each set, and points shaped by the overall change (activated vs. suppressed) (Fig. 3B).

Both the scatterplot of differentially expressed gene sets and the volcano plot were rendered using the ggplot2 v3.4.2 R package (94).

### Pathway Analysis

KEGG pathway analysis (42, 56) was performed using Pathways dre:00900, dre:00100, and dre:00140. These pathways were then colorized by relative gene expression using the pathview v1.38.0 R package (95) and overlaid on a manually redrawn pathway figure, combining the three individual pathways and showing only genes and portions of the pathways (genes and compounds) relevant to this study and species. Color bars on each gene node represent the relative gene expression of each sample in this study, scaled between [-1,1]. The top color bar of a given gene node represents the 10 wild-type samples and the bottom color bar represents the 10 mutant samples (Fig. 4).

### Statistics

All graph generation and statistical analyses, including calculation of means, and SEM, were performed using GraphPad Prism software (www.graphpad.com). D’Agostino and Pearson normality test was used to test whether data were normally distributed. If data were normally distributed, significance was assessed using t-tests with Welch’s correction or ANOVA with Dunnet’s multiple comparisons test. If data were not normally distributed, Mann-Whitney test or Kruskall-Wallis test with Dunn’s multiple comparisons test was used. *p*-values below 0.05 were considered statistically significant.

## Author Contributions

**Nicholas J. Santistevan:** Conceptualization, Methodology, Validation, Formal Analysis, Investigation, Data Curation, Visualization, Writing - Original Draft, Writing - Reviewing and Editing, Funding Acquisition. **Colby T. Ford:** Software, Formal Analysis, Resources, Data Curation, Visualization, Writing - Reviewing and Editing. **Cole S. Gilsdorf:** Investigation, Writing - Reviewing and Editing. **Yevgenya Grinblat:** Writing - Original Draft, Writing - Reviewing and Editing, Supervision, Project Administration, Funding Acquisition.

## Acknowledgements

We thank the members of the Grinblat lab for technical support and zebrafish line maintenance, and members of the labs of Dr. Mary Halloran, Dr. Katie Drerup, and Dr. David Ehrlich for help with zebrafish line maintenance. We thank Dr. Cecilia Moens for providing the *mecp2^Q63X^* mutant zebrafish line and Dr. Xinyu Zhao for expert insight and valuable advice. This work was funded by grants from the University of Wisconsin-Madison (Science and Medicine Graduate Research fellowship to N.J.S., Department of Integrative Biology funding to N.J.S, and the Office of the Vice Chancellor for Research and Graduate Education/Wisconsin Alumni Research Foundation to N.J.S. and Y. G), and by the National Science Foundation Graduate Research Fellowship Program under Grant Numbers DGE-1256259 and DGE-1747503 to N.J.S. Any opinions, findings, and conclusions or recommendations expressed in this material are those of the authors and do not necessarily reflect the views of the National Science Foundation.

## Data availability statement

All code, data, results, and additional analyses are openly available on GitHub at: https://github.com/colbyford/zebrafish_mecp2_expression. These data include RNA-seq gene expression values, gene set enrichment results, figure generation and statistics code, and additional pathway analyses.

## Ethics approval statement

All experiments and animal protocols were approved and conducted in accordance with the University of Wisconsin Institutional Animal Care and Use Committee (IACUC).

## Patient consent statement

Not applicable

## Permission to reproduce material from other sources

Not applicable

## Clinical trial registration

Not applicable

